# Detecting domain-level organization in genome-wide association study summary statistics using frequency–impact–reliability profiles

**DOI:** 10.64898/2026.07.01.735858

**Authors:** Hongdong Hao, Dian Chen, Xinyu Zhang, Hui Xue, Tianyu Meng

## Abstract

Genome-wide association studies (GWAS) identify trait-associated variants but are typically interpreted through single-variant significance and locus-level peaks, treating summary statistics as collections of independent signals. Here we show that GWAS summary statistics exhibit previously unrecognized local genetic–statistical organization along genomic coordinates.

We develop FIR-GWAS, a framework that integrates allele frequency, effect magnitude and statistical reliability to define frequency–impact–reliability (FIR) profiles and quantify their spatial continuity. Across EUR height GWAS, ancestry-specific datasets and eight additional human complex traits, we find consistent enrichment of same-profile adjacency and coordinate-contiguous FIR domains beyond chromosome-preserving null expectations. These patterns persist after removal of genome-wide significant variants and are reproducible across SNP- and window-based analyses.

We further show that FIR-domain architecture separates genome-wide significant structured regions from isolated association peaks and identifies subthreshold domains with coherent statistical organization. FIR-domain structure is consistently associated with regulatory annotations and trait-related gene sets, and highlights biologically plausible subthreshold candidate regions.

Across Arabidopsis and chicken GWAS, we observe that FIR-domain architecture is not restricted to human traits but recurs across independent association-summary landscapes. Decomposition analyses suggest that this spatial regularity arises from coordinated local continuity in allele frequency, effect size and statistical reliability.

Together, these results reveal GWAS summary statistics as structured genetic–statistical landscapes rather than collections of independent signals, defining a domain-level layer of organization that complements conventional single-variant and locus-based interpretation.

## 1 Introduction

Genome-wide association studies have identified large numbers of trait-associated variants and loci across human complex traits, providing a foundation for genetic discovery and downstream biological interpretation^1^. Most GWAS interpretation remains organized around single-variant significance, lead SNPs and locus-level association peaks^2^. This strategy has been effective for discovering robust trait-associated regions, but it treats GWAS summary statistics primarily as a collection of independent variant-level signals^2^. In practice, GWAS summary statistics are ordered along genomic coordinates, and each variant carries multiple statistical attributes beyond its P value, including allele frequency, effect estimate and standard error^3^. This raises the possibility that GWAS summary statistics may contain local spatial organization that is not captured by conventional locus-based interpretation^4^.

Although GWAS results are inherently coordinate-ordered, the spatial organization of their statistical attributes has not been systematically treated as a primary object of analysis^5^. Existing coordinate-aware approaches focus on locus definition, linkage disequilibrium structure, fine-mapping and functional annotation of associated regions^4,5^. These approaches are essential for translating association signals into candidate variants and genes, but they generally begin from significance-defined loci or predefined genomic features^2^. They do not directly examine whether neighbouring SNPs exhibit reproducible local similarity in allele frequency, effect magnitude or statistical reliability prior to downstream mapping^6^. As a result, a potential layer of organization between individual SNP-level signals and biological interpretation remains largely unexplored.

Allele frequency, effect magnitude and statistical reliability provide complementary views of GWAS summary statistics^3,6^. Allele frequency captures population context, effect magnitude reflects the estimated association strength, and statistical reliability reflects estimation stability relative to standard error^3^. Together, these dimensions define a local genetic-statistical state that cannot be reduced to P value alone^4^. Variants with similar association strength may differ substantially in frequency, effect size or reliability, whereas sub-threshold variants may still share coordinated local statistical profiles^6^. This suggests that integrating these dimensions may reveal latent structure in GWAS summary statistics that is not visible from single-variant significance alone^4^.

We therefore hypothesized that GWAS summary statistics contain non-random local frequency–impact–reliability architecture beyond individual association peaks^6^. Specifically, we proposed that neighbouring SNPs with similar allele frequency, effect magnitude and statistical reliability profiles exhibit enriched same-profile continuity along genomic coordinates and form coordinate-contiguous domains more frequently than expected under coordinate-preserving null models^5,7^. To test this hypothesis, we developed FIR-GWAS, a summary-statistic framework that assigns SNPs to frequency–impact–reliability profiles, quantifies local continuity and converts recurrent profile organization into permutation-scored FIR-defined domains^3,4^.

We applied FIR-GWAS to EUR height GWAS as a discovery setting and evaluated its generality across ancestry-specific height datasets and eight additional human complex traits^8,9^. We further developed FIR8 domain calling to distinguish genome-wide significant structured domains, genome-wide significant non-structured domains and non-GWS subthreshold structured domains^4^. To assess robustness, we evaluated alternative thresholds, null models, domain definitions, regulatory annotation and height-related gene context^10,11^. Finally, we extended the framework to Arabidopsis and chicken association datasets and decomposed the genetic-statistical basis of FIR spatial regularity^12,13^. Together, these analyses tested whether FIR-domain architecture represents a reproducible, scoreable and interpretable layer of local genetic-statistical organization in GWAS summary statistics, complementary to conventional lead-SNP and locus-based interpretation^2^.

## 2 Methods

### 2.1 Study design, data sources and quality control

We developed FIR-GWAS to characterize local spatial organization in GWAS summary statistics using three complementary dimensions: allele frequency, effect magnitude and statistical reliability^14^. The primary discovery analysis used EUR height GWAS summary statistics, and ancestry-specific height GWAS datasets were used for cross-ancestry validation. Cross-trait validation included eight additional human complex traits: BMI, LDL, ApoB, TG, uric acid, SBP, FEV1 and smoking. We further extended the framework to plant and animal GWAS datasets and examined the genetic-statistical basis of FIR spatial regularity in EUR height GWAS.

Human height GWAS summary statistics were obtained from the GIANT Consortium Yengo 2022 height GWAS resource, including EUR as the primary discovery dataset and AA, EAS, HIS and SAS datasets for cross-ancestry validation. Cross-trait human GWAS summary statistics were obtained from publicly available GWAS summary-statistic resources and included BMI (ieu-a-2), FEV1 (ukb-b-19657), LDL cholesterol (ieu-b-110), apolipoprotein B (ieu-b-108), triglycerides (ieu-b-111), serum uric acid (GCST90018977), systolic blood pressure (ukb-b-20175) and smoking initiation (ieu-b-4877). Cross-species validation used AraGWAS flowering-time association datasets for Arabidopsis thaliana, including FT10 (phenotype ID 261; AraGWAS:261; DOI: 10.21958/phenotype:261) and FT16 (phenotype ID 262; AraGWAS:262; DOI: 10.21958/phenotype:262), and Gallus gallus GWAS summary statistics for BW8, GW and EW from the Chicken AIL population GWAS summary data deposited in Zenodo record 15370428^12^. Gene and regulatory annotations were obtained from GENCODE v19, ENCODE cCRE hg19 and Roadmap 18-state chromatin annotations.

Human GWAS summary statistics were read from tabular or VCF-format files^3^. Required information included chromosome, position, SNP identifier, allele frequency, effect estimate, standard error and association statistic. For VCF datasets, effect size, standard error, LP or P value, allele frequency and available sample-size information were extracted from FORMAT or INFO fields when available. Only autosomal variants were retained for human analyses^15^. Variants were excluded if they had invalid chromosome or position, allele frequency outside the open interval (0,1), non-positive standard error, invalid P values or non-finite association statistics^3^.

Allele frequency was converted to minor allele frequency as MAF = min(AF, 1 − AF). Effect magnitude was defined as the absolute effect estimate, and statistical reliability was represented by |Z|, where Z = effect estimate / standard error^6^. Association strength was represented as LP = −log10(P); when LP was provided directly, P was derived as 10^−LP.

For the main human FIR analyses, association-enriched SNP sets were defined using LP > 4. The no-GWS sensitivity set excluded variants exceeding the conventional genome-wide significance threshold, defined as P < 5 × 10^−8 or equivalently LP > −log10(5 × 10^−8). Thus, variants retained in the no-GWS set satisfied P ≥ 5 × 10^−8 or LP ≤ −log10(5 × 10^−8). In the initial SNP-level and window-level analyses, an additional sensitivity set excluded the top 1% of LP values within each chromosome. For ancestry-specific height analyses, FIR cutoffs were calculated within each ancestry and analysis set. EUR-anchored and common-SNP sensitivity analyses were also performed to evaluate ancestry-dependent profile composition and shared-SNP robustness.

### 2.2 FIR profiling and local continuity analysis

Within each analysis set, SNPs were assigned frequency–impact–reliability profiles using MAF, absolute effect size and |Z|. High-frequency SNPs were defined by MAF greater than or equal to the within-dataset median. High-impact SNPs were defined by absolute effect size greater than or equal to the 75th percentile. High-reliability SNPs were defined by |Z| greater than or equal to the 75th percentile. FIR4 profiles combined frequency and impact groups, whereas FIR8 profiles additionally incorporated reliability^4^. FIR cutoffs were recalculated within each dataset, trait, ancestry or species-specific analysis set to avoid imposing a common scale across heterogeneous GWAS datasets.

As an auxiliary analysis, residual association strength was calculated by fitting LP against log-transformed absolute effect size, standard error and MAF. SNPs were classified into high or low residual LP groups using the 75th percentile of the residual distribution. The principal analyses focused on FIR4 and FIR8.

SNP-level FIR continuity was quantified after sorting variants by chromosome and position. Adjacent SNP pairs were defined as consecutive variants located on the same chromosome. The observed same-profile adjacency rate was the fraction of adjacent pairs sharing the same FIR profile. In the initial continuity analyses, profile labels were shuffled across analysed SNPs while preserving coordinate order, and the same-chromosome adjacent-pair structure was re-evaluated^16^. This procedure was repeated 1,000 times. Enrichment was calculated as the observed same-profile rate divided by the mean permuted rate, and empirical P values were calculated with a plus-one correction:

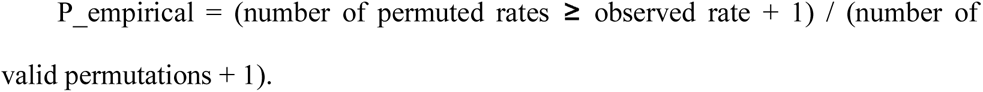

Window-level continuity was analysed by assigning variants to fixed genomic windows. In the initial analyses, 500 kb and 1 Mb windows were used. Within each occupied window, the dominant FIR profile was defined as the most frequent profile. Adjacent occupied windows on the same chromosome were tested for sharing the same dominant profile. Dominant-profile labels were permuted across occupied windows 1,000 times while preserving genomic order and same-chromosome window-pair structure.

The same FIR profiling, SNP-level continuity and window-level continuity procedures were applied independently to the eight human complex traits. Compact cross-trait signatures were generated from FIR4 and FIR8 SNP-level enrichment ratios and 500 kb window-level enrichment ratios.

### 2.3 FIR-defined domain architecture and domain calling

FIR domains were defined as coordinate-contiguous same-profile SNP runs^17^. A new domain was initiated whenever the chromosome changed or the FIR profile differed from that of the preceding SNP. This definition does not rely on predefined genomic intervals; domains were inferred directly from the spatial continuity of FIR profiles.

Formal FIR-domain architecture analysis was first performed in EUR height GWAS and then extended to ancestry-specific height GWAS datasets and eight human complex traits. For each dataset, the main LP > 4 set and the no-GWS set were analysed. Observed same-profile blocks were summarized using length- and SNP-count-based metrics, including high-percentile block length, high-percentile SNPs per block and N50 metrics. To generate null expectations, FIR profile labels were shuffled within chromosomes while preserving SNP coordinates and chromosome-specific SNP density. Block metrics were recomputed across 500 permutations. Enrichment was calculated as the observed value divided by the mean permuted value, and log2 enrichment was calculated as log2(observed/permutation mean).

A compact architecture score was calculated from five core metrics: q95 block length, q99 block length, q95 SNPs per block, q99 SNPs per block and N50 SNPs per block. The score was defined as the mean log2 enrichment across these metrics. Scale-dependent continuity was further assessed using fixed windows of 100 kb, 250 kb, 500 kb, 1 Mb and 2 Mb, with dominant-profile labels permuted within chromosomes 500 times.

After architecture-level testing, FIR8 domain calling was performed in the EUR height GWAS LP > 4 SNP set. FIR8 domains were called as consecutive same-FIR8 SNP runs. For each observed domain, we recorded genomic coordinates, domain length, SNP count, FIR8 profile, frequency group, impact group, reliability group, summary FIR variables, lead SNP, lead P value, lead LP and genome-wide significance status.

Each FIR8 domain was assigned a statistical architecture class according to its impact and reliability status. High-impact/high-reliability domains were labelled signal-enriched, high-impact/low-reliability domains impact-only, low-impact/high-reliability domains reliability-dominant and low-impact/low-reliability domains background. These classes described the statistical character of each domain; structured-domain calling was based on permutation-derived domain length and SNP-count percentiles.

To score domain structure, FIR8 profile labels were shuffled within chromosomes 1,000 times, and domains were re-called from each permuted profile assignment. For each observed domain, length and SNP count were compared with same-profile permuted domains. The length percentile was defined as the proportion of same-profile permuted domains with length less than or equal to the observed length, and the SNP-count percentile was defined analogously. The domain structure score was calculated as the mean of these two percentiles.

Domains were considered raw structured at q90, q95 or q99 if either the length percentile or SNP-count percentile reached the corresponding threshold. Final structured labels additionally required at least two SNPs. The q95 final structured label was used as the main structured-domain definition. Domains passing q99 were considered very-high-confidence structured, domains passing q95 high-confidence structured and domains passing q90 moderate structured.

Domains were prioritized by combining genome-wide significance status, q95 final structure status and maximum LP. Domains containing at least one genome-wide significant SNP and passing q95 structure criteria were classified as GWS structured domains. Domains containing a genome-wide significant SNP but failing q95 structure criteria were classified as GWS non-structured domains. Non-GWS domains passing q95 structure criteria and having maximum LP ≥ 6 were classified as non-GWS subthreshold structured domains. Non-GWS domains passing q95 structure criteria but having maximum LP < 6 were classified as non-GWS background structured domains. A higher subthreshold tier, maximum LP ≥ 7, was recorded, but the main non-GWS subthreshold structured class used maximum LP ≥ 6.

### 2.4 Robustness and sensitivity analyses

Robustness analyses evaluated whether FIR-domain calling depended on the LP input threshold, null model, structured-domain definition, genomic region inclusion or FIR splitting strategy. In each setting, domains were re-called after re-filtering the SNP set, reassigning FIR8 profiles and re-estimating the appropriate null background. Structured-domain enrichment was summarized as the observed number of structured domains divided by the mean number expected under the null model, with empirical upper-tail P values calculated using a plus-one correction.

LP-threshold robustness repeated domain calling at LP ≥ 3, 3.5, 4, 4.5 and 5 using the median FIR split, chromosome-shuffle null model, q95 combined structure criterion and a minimum of two SNPs per structured domain. Each threshold used 100 permutations. Null-model robustness compared genome-wide shuffling, chromosome-level shuffling, 1 Mb local-window shuffling and circular shifting of FIR8 labels within chromosomes, using LP ≥ 4 and 100 permutations.

Parameter robustness evaluated 27 structured-domain definitions formed by combining three percentile thresholds, q90, q95 and q99; three minimum-SNP requirements, at least 2, 3 or 5 SNPs; and three structural metrics, combined, length-only and SNP-count-only. Leave-region-out robustness repeated the analysis after removing each autosome individually and after removing the top 1% of domains ranked by maximum LP, domain length or SNP count, using 50 permutations per setting. FIR-split robustness tested median, q40/60, tertile-high and quartile-high split strategies, using 100 permutations per setting.

### 2.5 Biological annotation and candidate-domain prioritization

The EUR height FIR8 domain catalogue was annotated using gene models, regulatory annotations and curated height-biology gene sets. This analysis used the same FIR-domain calling framework, including LP > 4 SNP selection, FIR8 assignment, chromosome-stratified profile permutations, q95 final structured-domain classification and priority groups defined by genome-wide significance and structure status.

Gene annotations were obtained from GENCODE v19^18^. Gene records were restricted to autosomes and parsed for gene identifier, gene name, gene type and strand. Promoter intervals were defined as 2 kb upstream and 500 bp downstream of the transcription start site, with strand-specific orientation. Domains were mapped to genes by direct gene-body overlap, presence of genes within 100 kb and nearest-gene distance.

Regulatory annotations included GENCODE promoter intervals, ENCODE cCRE annotations and Roadmap 18-state chromatin annotations^10,11^. ENCODE cCRE features were grouped into promoter-like, proximal enhancer-like, distal enhancer-like and accessible-chromatin categories. Roadmap states were summarized into active TSS, TSS-flanking, active enhancer, weak enhancer, genic enhancer and transcribed groups. A fixed skeletal-growth core Roadmap epigenome group was used for the core regulatory annotation analysis, with an extended annotation set generated for supplementary analyses.

Domain–feature overlaps were calculated by genomic interval intersection^10^. For each biological domain class, feature enrichment was evaluated using Fisher’s exact test by comparing feature-overlapping and non-overlapping domains against the remaining domain background. P values were adjusted for multiple testing using false-discovery rate correction.

Curated height-biology gene sets included skeletal-growth genes, WNT/BMP/TGF/FGF/HH developmental signalling genes, growth-plate and chondrocyte genes, extracellular-matrix and collagen genes, and endocrine growth-axis genes. Domain-level gene mappings were intersected with these sets, and enrichment was evaluated using domain-level gene-set overlap statistics with false-discovery rate correction.

Non-GWS subthreshold structured domains were prioritized as candidate domains. For each candidate domain, nearest gene, nearest-gene distance, overlapping genes, genes within 100 kb, height-biology genes, regulatory feature groups and regulatory overlap counts were recorded. A biology priority score combined standardized domain structure score, standardized maximum LP, standardized log-transformed SNP count, height-biology gene overlap and regulatory overlap. Candidate domains were ranked by this score. These annotations were used for candidate prioritization only and were not interpreted as functional validation.

### 2.6 Cross-species validation and mechanistic decomposition

To evaluate whether FIR-domain architecture recurred beyond human GWAS, the framework was applied to plant and animal association datasets. Cross-species validation included two Arabidopsis thaliana flowering-time traits, FT16 and FT10, and three Gallus gallus growth or production traits, BW8, GW and EW. Arabidopsis association results were read from AraGWAS HDF5 files, and chicken GWAS datasets were read from tab-delimited summary-statistic files. Across species, variants were standardized to chromosome, position, SNP identifier, MAF, effect estimate, standard error, P value, LP, absolute effect estimate and |Z|.

Within each cross-species trait, association-enriched variant sets were defined by trait-specific top fractions of the LP distribution. The analysed fractions were the top 5%, top 10% and top 20% of variants. FIR8 profiles were constructed separately within each trait and top-fraction dataset using median MAF, the 75th percentile of absolute effect estimate and the 75th percentile of |Z|. Same-profile adjacency and FIR-domain architecture were quantified using the same coordinate-based definitions as in the human analyses. FIR8 labels were permuted within chromosomes 1,000 times to generate null expectations. The primary analysis focused on the Top10 dataset, with Top5 and Top20 used as sensitivity analyses. Cross-species recurrence was interpreted as recurrence of FIR-domain statistical architecture across independent association-summary landscapes, not as evidence for shared biological mechanisms.

To investigate the origin of FIR spatial regularity, a mechanistic decomposition analysis was performed in EUR height GWAS using the top 10% of variants by LP. FIR profiles were constructed from MAF, |BETA| and |Z| using median MAF and the 75th percentiles of |BETA| and |Z|. Lower-dimensional profiles were also defined for frequency, impact, reliability and their pairwise combinations.

Adjacent same-chromosome SNP pairs were used to evaluate local continuity in the continuous FIR components. For each pair, physical distance and absolute differences in MAF, |BETA|, |Z|, LP and SE were calculated. Component-level spatial autocorrelation was tested by comparing adjacent-pair deltas with chromosome-matched random-pair deltas sampled in proportion to chromosome-specific SNP counts, using 1,000 permutations. The contribution of frequency, impact and reliability to same-profile adjacency was assessed by independently permuting frequency, impact and reliability bins within chromosomes 1,000 times and reconstructing individual, pairwise and full FIR profiles.

Distance-stratified FIR8 enrichment was assessed by grouping adjacent pairs by physical distance and comparing same-FIR8 rates with chromosome-preserving permutation backgrounds. Within-domain homogeneity and F/I/R coupling were evaluated by comparing observed FIR-domain segments with random matched segments, focusing on within-domain variance in MAF, |BETA| and |Z| and correlations among the three FIR components. Boundary transitions were analysed using 50 SNPs on each side of FIR-domain boundaries and compared with a random boundary background using 500 permutations. Finally, logistic regression was used to model same-FIR8 adjacency as a function of physical distance, allele-frequency variables, uncertainty variables based on SE, effect-magnitude variables based on |BETA|, and reliability/significance variables based on |Z| and LP. Model fit was summarized using McFadden’s pseudo-R²^19^.

## 3 Results

### 3.1 FIR-GWAS identifies local profile continuity in height GWAS

We first asked whether GWAS summary statistics contain local spatial organization beyond individual association peaks. To address this, we developed a frequency–impact–reliability (FIR) profiling framework, in which each SNP was represented by allele frequency, effect magnitude and statistical reliability. These features were combined into FIR4 or FIR8 profiles and examined along genomic coordinates for same-profile continuity (Fig. 1A).

**Figure 1.**
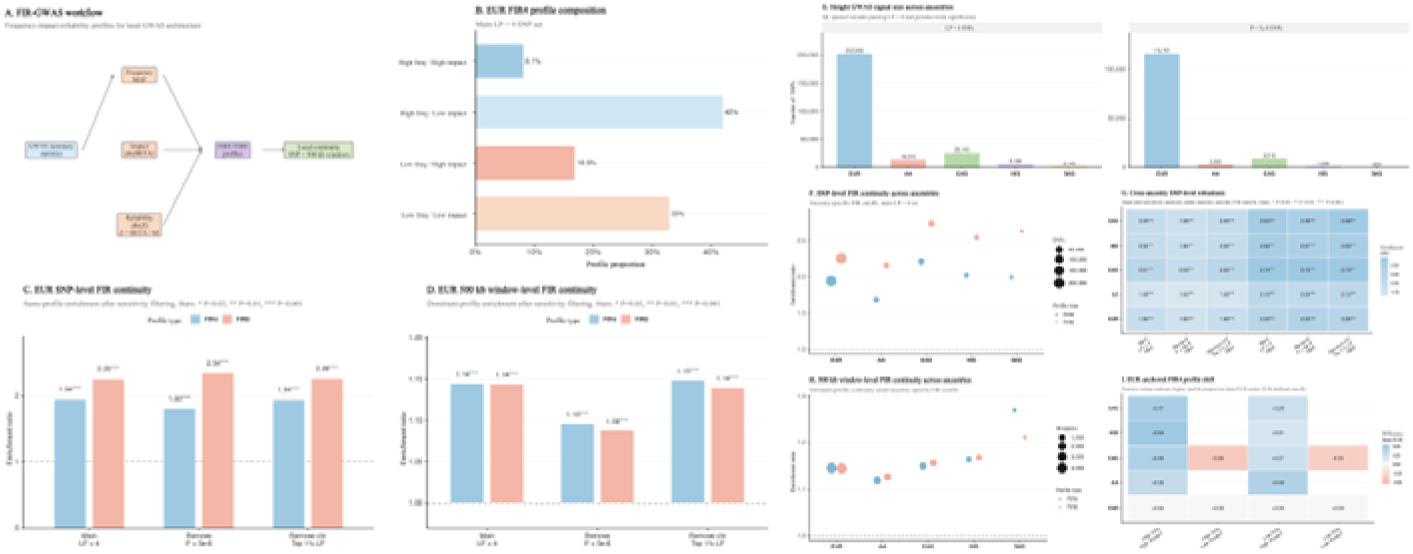
FIR-GWAS framework and EUR height discovery. A, FIR-GWAS workflow converting GWAS summary statistics into FIR profiles (MAF, |β|, |Z|) and genomic domains. B, FIR4 profile distribution in EUR height GWAS (LP > 4). C, SNP-level FIR continuity showing same-profile enrichment. D, 500 kb window-level FIR continuity. E, height GWAS signal size across ancestries. F, SNP-level FIR continuity across ancestries. G, cross-ancestry enrichment heatmap. H, window-level continuity across ancestries. I, EUR-anchored profile shifts

In the EUR height GWAS LP > 4 SNP set, FIR4 profiles were unevenly distributed. High-frequency/low-impact and low-frequency/low-impact SNPs accounted for 42.05% and 32.95% of variants, respectively, whereas low-frequency/high-impact and high-frequency/high-impact SNPs accounted for 16.85% and 8.15% (Fig. 1B). Despite this compositional imbalance, adjacent SNPs were substantially more likely to share the same FIR profile than expected under chromosome-stratified permutation. In the main EUR analysis, FIR4 and FIR8 same-profile adjacency showed enrichment ratios of 1.94 and 2.25, respectively, both with empirical P = 9.99 × 10^-4 (Fig. 1C). This enrichment persisted after removing genome-wide significant SNPs and after removing the top 1% LP SNPs within each chromosome, indicating that local FIR continuity was not driven solely by classical association peaks or extreme signals.

The same pattern extended to a broader genomic scale. In 500 kb windows, dominant-profile continuity remained enriched for both FIR4 and FIR8 in the EUR dataset, with enrichment ratios around 1.14 in the main analysis and comparable enrichment after sensitivity filtering (Fig. 1D). We then applied the framework to ancestry-specific height GWAS datasets. Although the number of LP > 4 SNPs varied markedly across ancestries, from 202,996 in EUR to 2,143 in SAS, same-profile enrichment was consistently observed. Across ancestries, SNP-level FIR4 enrichment ranged from 1.69 to 2.21 and FIR8 enrichment from 2.16 to 2.74, with empirical P = 9.99 × 10^-4 across all ancestry datasets (Fig. 1E–G). Window-level continuity also remained directionally consistent, with 500 kb FIR4 and FIR8 enrichment observed across ancestry datasets (Fig. 1H). EUR-anchored FIR4 profiling further showed that non-EUR ancestries differed in profile composition, particularly through higher proportions of high-impact profiles, indicating that recurrent FIR continuity was not simply a consequence of identical profile distributions across ancestries (Fig. 1I).

Thus, FIR profiling converts height GWAS summary statistics into local statistical profiles and reveals recurrent SNP-level and window-level spatial continuity across ancestry-specific datasets.

### 3.2 FIR continuity generalizes across human complex traits

We next tested whether FIR continuity was specific to height or reflected a broader property of human complex-trait GWAS. We analysed eight additional traits spanning anthropometric, lipid, metabolic, vascular, pulmonary and behavioural systems: BMI, LDL, ApoB, TG, uric acid, SBP, FEV1 and smoking. The number of LP > 4 SNPs varied widely across traits, from 9,251 in BMI to 191,365 in TG, and genome-wide significant SNP counts also differed substantially (Fig. 2A). This variation provided a broad setting in which to assess whether FIR continuity depended on the size or strength of the association signal set.

**Figure 2.**
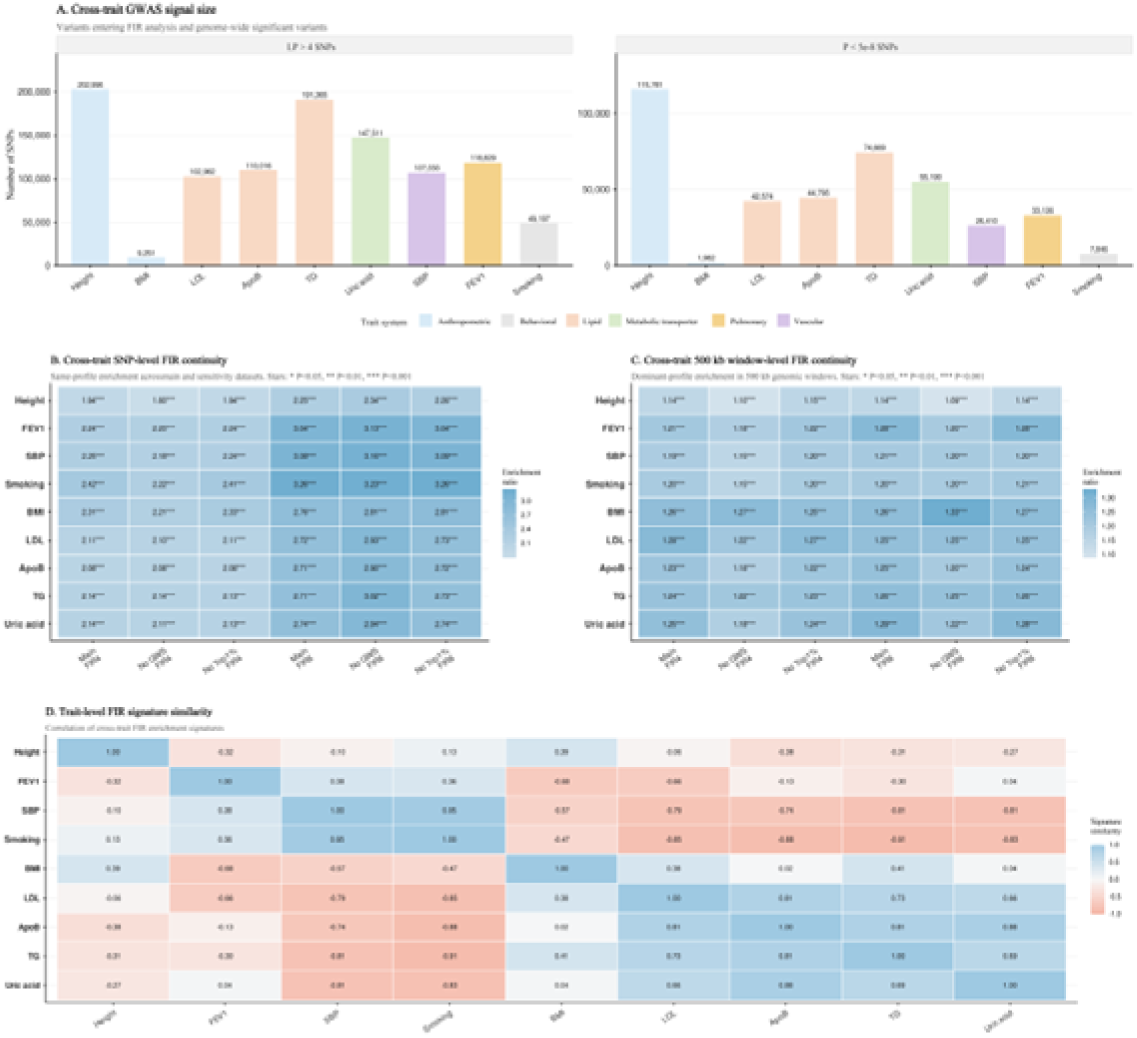
FIR continuity across human complex traits. A, Cross-trait GWAS signal size, showing the numbers of LP > 4 SNPs and genome-wide significant variants across height and eight additional traits. B, SNP-level FIR continuity across traits, showing consistent same-profile enrichment for FIR4 and FIR8 in the main, no-GWS and top-1%-removed analyses. C, 500 kb window-level FIR continuity across traits, showing reproducible dominant-profile enrichment at broader genomic scale. D, Trait-level FIR signature similarity, showing shared but non-identical FIR patterns across phenotypes, with stronger similarity among related traits.

Across all eight non-height traits, SNP-level same-profile enrichment was consistently detected. In the main LP > 4 analysis, FIR4 enrichment ranged from 2.08 to 2.42, whereas FIR8 enrichment was stronger, ranging from 2.71 to 3.26, with empirical P = 9.99 × 10^-4 for all traits (Fig. 2B). Enrichment remained after excluding genome-wide significant SNPs and after removing the top 1% LP SNPs per chromosome, supporting that cross-trait FIR continuity was not attributable to a small number of extreme association peaks.

The signal also generalized to 500 kb window-level continuity. Across the eight traits, FIR4 window-level enrichment ranged from 1.19 to 1.28 and FIR8 enrichment from 1.20 to 1.29 in the main analysis, again with empirical P = 9.99 × 10^-4 across traits (Fig. 2C). Although these window-level ratios were smaller than SNP-level enrichment ratios, their consistent direction across traits indicates that FIR continuity extends beyond immediate SNP adjacency.

Trait-level FIR signatures were not identical across phenotypes. Lipid-related traits showed high similarity among LDL, ApoB and TG, whereas smoking and SBP formed another strongly correlated pattern. Height showed weaker similarity to most non-height traits (Fig. 2D). Thus, FIR continuity is recurrent across complex-trait GWAS, whereas the detailed FIR signature remains trait-dependent.

### 3.3 FIR-defined genomic domains represent the core spatial architecture of GWAS summary statistics

The preceding analyses established local FIR continuity, but did not yet define a higher-order genomic structure. We therefore formalized FIR-defined genomic domains as contiguous same-profile SNP blocks along physical coordinates and tested whether these blocks were longer and denser than expected under chromosome-stratified profile permutation. This shifted the analysis from local continuity to domain-level architecture.

In Height EUR GWAS, FIR8 profiles produced a genome-wide domain-like landscape across hg19 chromosomes, with background, reliability-dominant, impact-only and signal-enriched architectures appearing as spatially organized segments rather than randomly scattered labels (Fig. 3A). A representative signal-enriched region further showed that SNP-level FIR8 classes, local association intensity and domain tracks aligned within a compact genomic interval (Fig. 3B).

**Figure 3.**
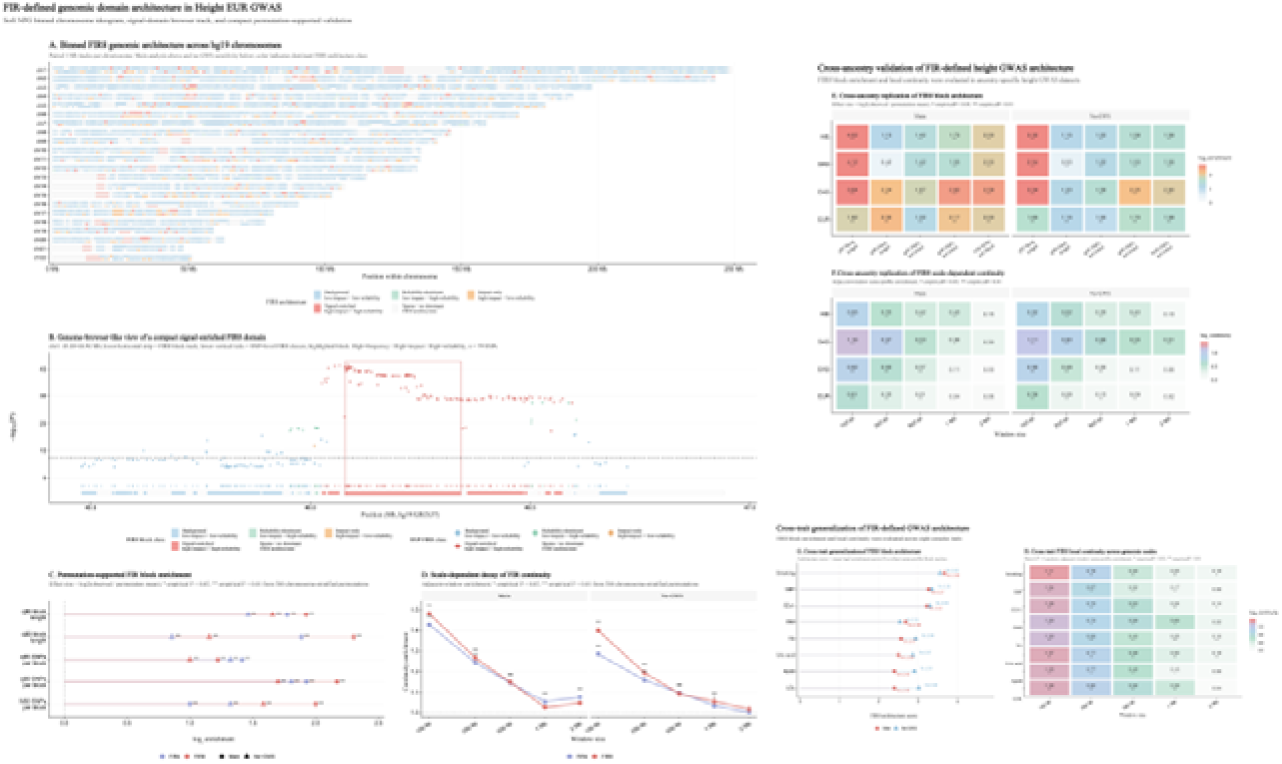
FIR-defined genomic domain architecture in EUR height GWAS and cross-ancestry validation. A, Genome-wide binned FIR8 architecture across hg19 chromosomes showing spatial distribution of FIR-defined blocks. B, Genome-browser view of a representative FIR-enriched domain with SNP-level signal and domain boundaries. C, Permutation-based enrichment of FIR block structure across multiple domain metrics (length, SNP count, N50). D, Scale-dependent decay of FIR continuity across increasing genomic window sizes. E, Cross-ancestry validation of FIR block enrichment in height GWAS across EUR, EAS, SAS and HIS datasets. F, Cross-ancestry replication of scale-dependent FIR continuity under main and no-GWS settings.

Permutation-supported block analysis confirmed that observed FIR domains exceeded null expectations. In the main Height EUR analysis, FIR4 q95 and q99 block-length enrichment reached 3.42 and 3.70, whereas FIR8 reached 3.79 and 4.93; SNP-per-block metrics were also enriched, including FIR8 q99 SNPs per block at 4.50. All tested core block metrics reached empirical P = 0.001996 (Fig. 3C). Importantly, domain-level enrichment persisted after removing genome-wide significant SNPs, indicating that FIR-defined architecture is not merely a restatement of conventional significant loci.

FIR-domain continuity was strongly scale-dependent. In the Height EUR main analysis, FIR8 window continuity decreased from 1.53 at 100 kb to 1.16 at 500 kb and approached the null at 1–2 Mb (Fig. 3D). The same decay pattern was observed after excluding genome-wide significant SNPs, placing the principal FIR-domain signal at local to intermediate genomic scales rather than at broad chromosome-wide structure.

This domain architecture extended beyond EUR height. In ancestry-specific height GWAS datasets, FIR8 architecture scores were positive across EUR, EAS, HIS and SAS, with main-analysis scores ranging from 1.90 to 2.59 (Fig. 3E). Multiscale continuity again showed strongest enrichment at 100–500 kb and attenuation at larger windows (Fig. 3F). Across eight human complex traits, FIR8 block architecture was also consistently detected, with main-analysis architecture scores ranging from 2.40 to 3.66 and all core block metrics significant across traits (Fig. 3G). Cross-trait window continuity likewise showed strongest enrichment at local scales (Fig. 3H).

Together, these analyses identify FIR-defined genomic domain architecture as the central spatial regularity underlying the earlier SNP- and window-level observations. Same-profile SNPs form non-random blocks, these blocks exceed permutation expectations, the architecture is strongest at local to intermediate scales, and the pattern recurs across ancestry-specific height GWAS and multiple human traits. This domain-level representation captures an organization of GWAS summary statistics that is not visible from single-SNP significance alone.

### 3.4 FIR-domain calling provides a robust domain-level framework

We next converted FIR-domain architecture into an explicit domain-calling framework. This framework transforms SNP-level FIR8 profiles into contiguous, structure-scored genomic domains using chromosome-stratified permutation backgrounds (Fig. 4A).

**Figure 4.**
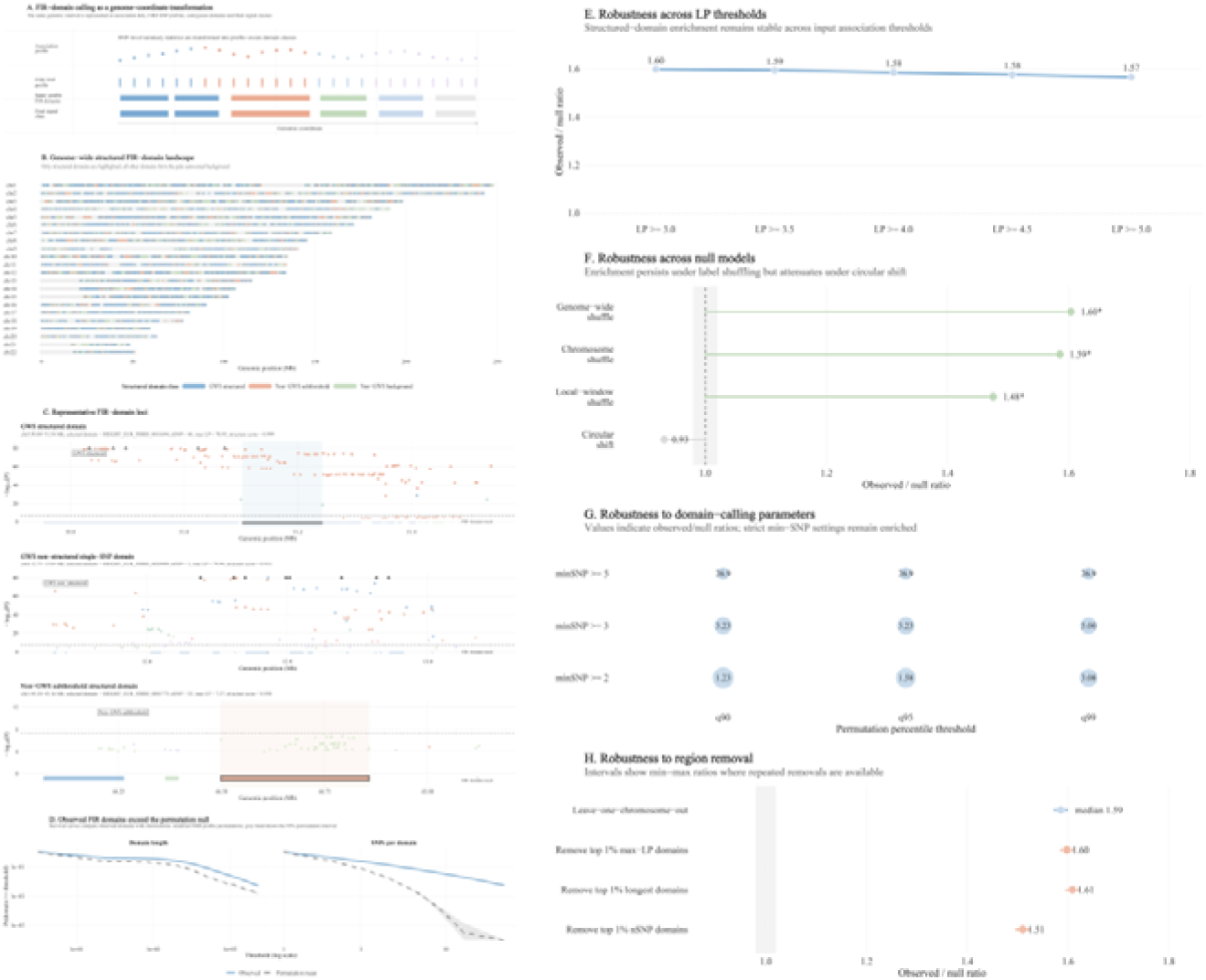
FIR-domain calling provides a robust domain-level framework for GWAS summary statistics. A, FIR-domain calling converts SNP-level FIR8 profiles into coordinate-contiguous genomic domains using chromosome-stratified permutation. B, Genome-wide FIR-domain classes in EUR height GWAS including GWS structured, GWS non-structured, non-GWS structured and non-GWS background domains. C, Representative loci showing GWS structured domains, GWS non-structured peaks and non-GWS structured domains. D, FIR-domain length and SNP-count distributions compared with permutation-derived null expectations. E, Robustness across LP thresholds. F, Robustness across null models including genome-wide, chromosome, local-window and circular shift permutations. G, Robustness across domain-calling parameters including percentile thresholds and minimum SNP requirements. H, Leave-region-out sensitivity analysis.

In Height EUR GWAS, FIR-domain calling identified 94,466 FIR8 domains, including 19,100 GWS structured domains, 38,830 GWS non-structured domains, 2,473 non-GWS subthreshold structured domains and 3,191 non-GWS background structured domains (Fig. 4B). The contrast between GWS structured and GWS non-structured domains was particularly informative. GWS structured domains had a median length of 7,953 bp and median SNP count of 3, whereas GWS non-structured domains had a median length of 1 bp and median SNP count of 1. Thus, genome-wide significance alone did not imply regional structure. Conversely, non-GWS subthreshold structured domains had a median length of 23,401 bp and median SNP count of 4, indicating that FIR-domain calling can identify structured regions below the conventional significance threshold.

Representative loci illustrated these distinctions: a GWS structured domain formed a compact multi-SNP association region, a GWS non-structured peak remained essentially isolated, and a non-GWS subthreshold structured domain showed regional structure without crossing the genome-wide significance threshold (Fig. 4C). Across all domains, observed domain-length and SNP-count distributions showed heavier tails than permutation backgrounds (Fig. 4D), supporting that the called domains reflect profile-aware genomic structure rather than random label runs.

Robustness analyses further supported the framework. Structured-domain enrichment was stable across LP thresholds from ≥3.0 to ≥5.0, with observed/null ratios from 1.57 to 1.60 (Fig. 4E). The signal persisted under genome-wide, chromosome and local-window shuffling, but attenuated under circular shift, consistent with the dependence of FIR architecture on the original local genomic arrangement (Fig. 4F). Parameter-grid analysis showed that enrichment remained under alternative percentile thresholds and minimum-SNP requirements, with 25 of 27 settings classified as strong and 2 as supportive (Fig. 4G). Leave-region-out tests showed that removing individual chromosomes or top extreme domains did not abolish the signal (Fig. 4H).

These results establish FIR-domain calling as a stable framework for converting GWAS summary statistics into structure-scored genomic domains and for separating isolated significant peaks from multi-SNP structured regions.

### 3.5 FIR domains connect statistical architecture to height-related regulatory biology

After establishing a stable domain-calling framework, we evaluated whether FIR-defined domains carry biological interpretability. We integrated domain classes with gene mapping, ENCODE cCRE annotations, Roadmap 18-state chromatin states and curated height-biology gene sets (Fig. 5A).

**Figure 5.**
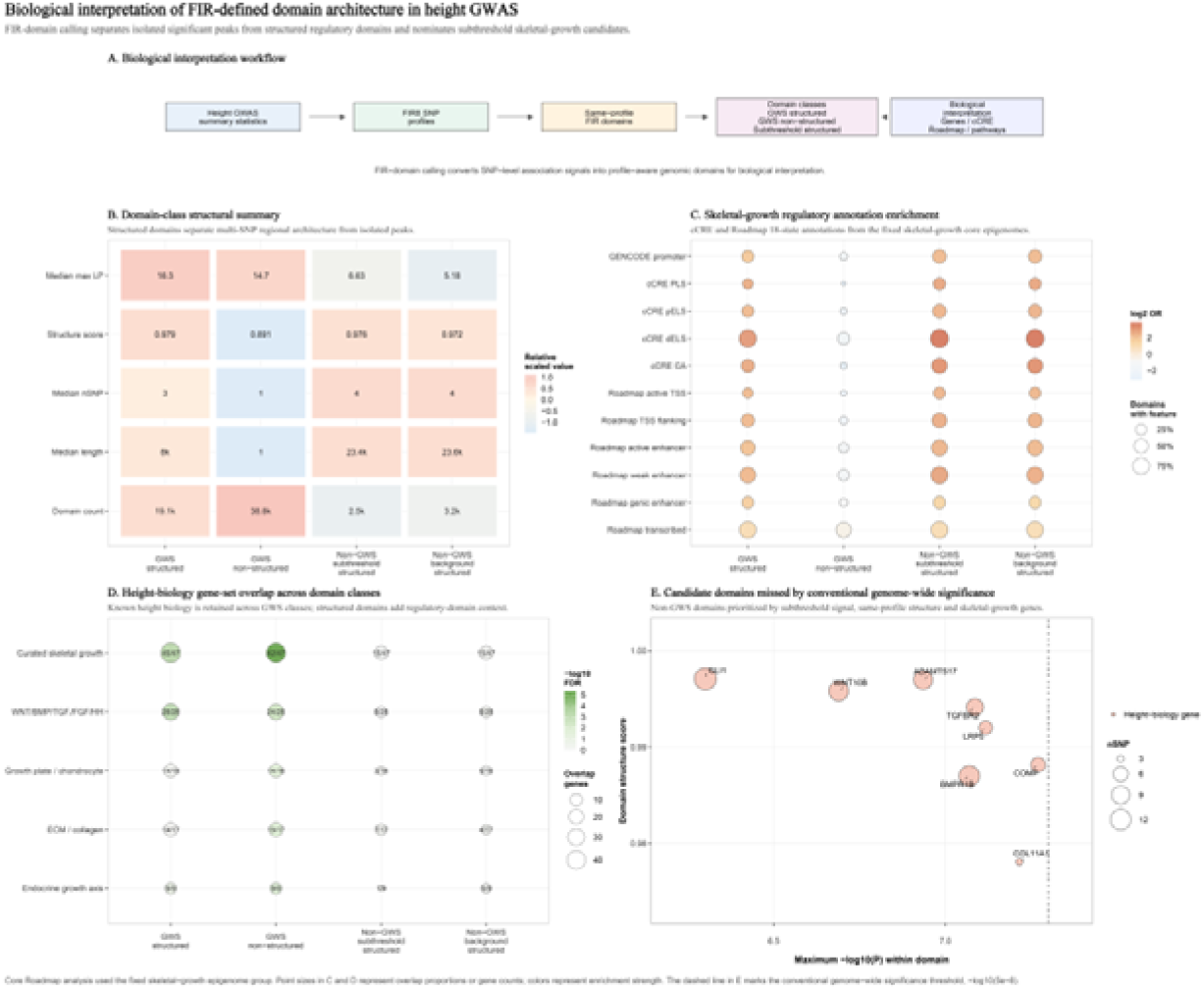
FIR domains connect statistical architecture to height-related regulatory biology. A, Overview of biological interpretation workflow linking FIR-domain classes to gene mapping, cCRE, Roadmap chromatin states and curated height-biology gene sets. B, Structural summary of domain classes showing differences in association strength, domain size and SNP content across GWS structured, GWS non-structured, non-GWS structured and non-GWS background domains. C, Regulatory annotation enrichment across domain classes using cCRE and Roadmap 18-state chromatin annotations. D, Height-biology gene-set overlap across domain classes for curated skeletal-growth and developmental signalling pathways. E, Candidate non-GWS structured domains highlighting subthreshold loci overlapping height-related genes including BMPR1B, ADAMTS17, TGFBR2, COMP, WNT10B, LRP5, GLI1 and COL11A1.

Different domain classes showed distinct structural and annotation profiles. GWS structured domains combined strong association signals with regional structure, whereas GWS non-structured domains were more numerous but had a median length of only 1 bp and a median SNP count of 1 (Fig. 5B). Non-GWS subthreshold structured domains were below the genome-wide significance threshold but showed high structure scores and multi-SNP regional organization, defining a candidate class distinct from simple background.

Structured domains were consistently more regulatory-annotation rich than non-structured domains. GWS structured domains were enriched for promoter-like, enhancer-like and skeletal-growth Roadmap annotations, including cCRE distal enhancer-like, accessible chromatin and promoter-like features (Fig. 5C). By contrast, GWS non-structured domains were depleted across many of these regulatory annotations, suggesting that some conventional significant peaks represent isolated statistical signals rather than structured regulatory domains. Non-GWS subthreshold structured domains also showed strong enrichment for cCRE enhancer-like, accessible chromatin and Roadmap enhancer/TSS-related annotations, indicating that subthreshold structured domains can carry regulatory context despite not reaching genome-wide significance.

Height-biology gene-set overlap further supported biological interpretability. GWS structured domains covered 45/47 curated skeletal-growth genes and all 28 WNT/BMP/TGF/FGF/HH developmental signalling genes, while GWS non-structured domains also retained many known height genes (Fig. 5D). Non-GWS structured domains showed weaker global enrichment but still contained subsets of skeletal-growth, developmental signalling, growth-plate, ECM/collagen and endocrine growth-axis genes. This pattern suggests that GWS classes capture much of the established height-biology signal, whereas non-GWS structured domains may prioritize additional candidate regions.

Focusing on non-GWS subthreshold structured domains, FIR-domain annotation nominated several biologically plausible candidate regions missed by conventional genome-wide significance. These included domains overlapping or neighbouring BMPR1B, ADAMTS17, TGFBR2, COMP, WNT10B, LRP5, GLI1 and COL11A1 (Fig. 5E). For example, BMPR1B, ADAMTS17, TGFBR2 and COMP candidate domains all remained below the conventional genome-wide significance threshold but showed high structure scores and multi-SNP regional support. These findings do not constitute functional validation, but they show that FIR-domain calling can connect subthreshold GWAS signal structure to regulatory annotation and height-related gene context.

### 3.6 FIR-domain architecture recurs across plant and animal GWAS

To test whether FIR-domain architecture is restricted to human GWAS, we extended the analysis to non-human association datasets. We analysed two Arabidopsis thaliana flowering-time traits and three Gallus gallus growth or production traits, applying trait-specific FIR8 profiling to the top 5%, 10% and 20% association variants and testing same-profile structure against chromosome-stratified permutations.

In the primary Top10 analysis, all plant and animal traits showed same-profile adjacency enrichment. Arabidopsis FT16 and FT10 had observed/null ratios of 1.65 and 1.66, whereas chicken BW8, GW and EW showed stronger enrichment of 2.76, 2.37 and 3.31, respectively, all with empirical P = 9.99 × 10^-4 (Fig. 6A). Domain-level metrics showed the same pattern. For q95 domain length, observed/null ratios were 1.58 for both Arabidopsis traits and ranged from 4.34 to 8.87 in chicken traits; for q95 SNPs per domain, ratios ranged from 1.33 in Arabidopsis to 5.63 in chicken EW (Fig. 6B).

**Figure 6.**
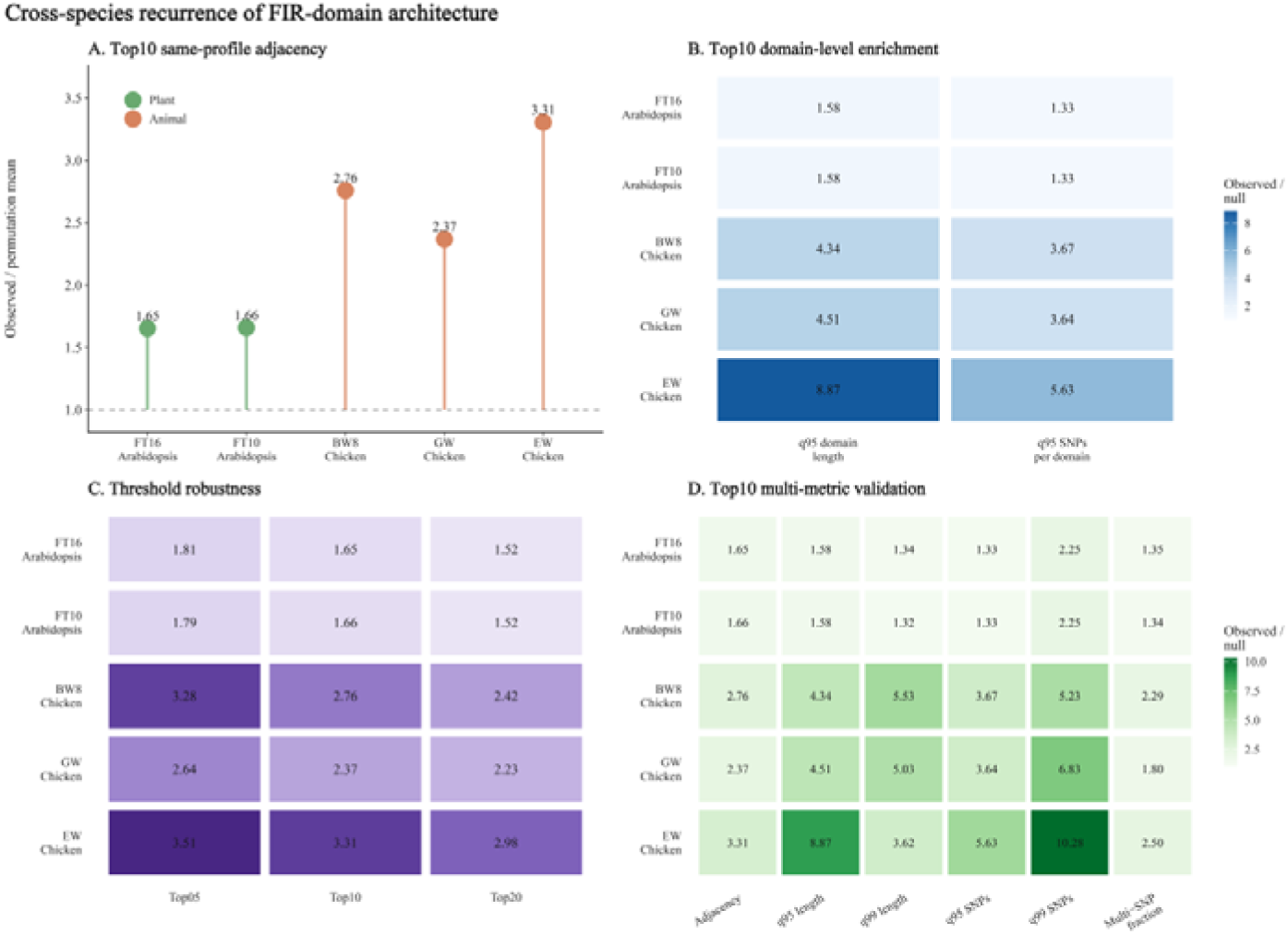
FIR-domain architecture recurs across plant and animal GWAS. A, Top10 same-profile FIR adjacency enrichment across Arabidopsis thaliana (FT16, FT10) and Gallus gallus (BW8, GW, EW). B, Top10 domain-level enrichment across species for q95 domain length and SNPs per domain. C, Threshold robustness across Top5, Top10 and Top20 variant sets. D, Multi-metric validation of FIR-domain architecture across adjacency, domain length, SNP count and multi-SNP fraction.

The recurrence was not specific to the Top10 threshold. Across Top5, Top10 and Top20 inputs, all five traits retained same-profile enrichment, although ratios generally declined as broader variant sets were included (Fig. 6C). Multi-metric validation showed enrichment for adjacency, high-percentile domain length, high-percentile SNPs per domain and multi-SNP domain fraction across all traits (Fig. 6D).

These cross-species analyses indicate that FIR-domain architecture is not confined to human height or human complex-trait GWAS. Rather, it recurs in independent plant and animal association-summary landscapes. This recurrence should be interpreted as evidence for a generalizable statistical architecture of GWAS summary data, not as evidence that different species or traits share identical biological mechanisms.

### 3.7 Local genetic-statistical continuity underlies FIR spatial regularity

Finally, we investigated why FIR spatial regularity emerges. We returned to the continuous variables underlying FIR profiles and tested whether local continuity, joint F/I/R coupling, distance persistence, domain homogeneity, boundary shifts and adjacent-pair modelling could account for the observed architecture (Fig. 7A).

**Figure 7.**
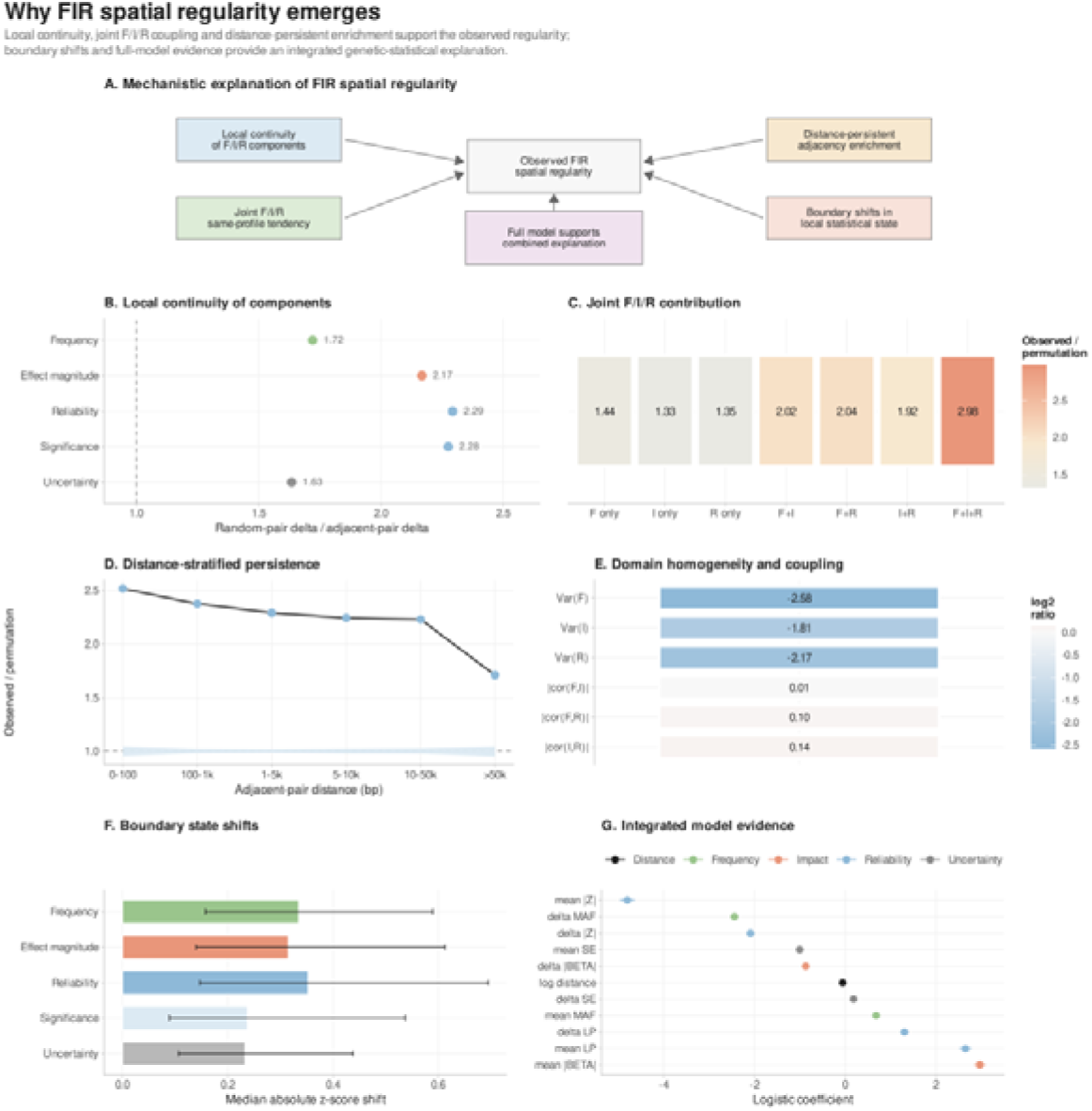
Local genetic-statistical continuity underlies FIR spatial regularity. A, Mechanistic framework for FIR spatial regularity integrating local continuity, joint F/I/R coupling, distance persistence, boundary shifts and full-model evidence. B, Local continuity of FIR components showing higher similarity in adjacent SNP pairs compared with random pairs across MAF, |β|, |Z|, LP and SE. C, Joint contribution of F/I/R components to same-profile adjacency, with strongest enrichment observed for full FIR8 profiles. D, Distance-stratified persistence of FIR continuity across increasing genomic distances. E, Domain homogeneity and coupling showing reduced within-domain variance in FIR components and weak pairwise correlations. F, Boundary state shifts across FIR-domain limits. G, Integrated model evidence from logistic regression showing combined contribution of distance, frequency, impact and reliability variables to same-FIR adjacency.

Adjacent SNPs were more similar than chromosome-matched random SNP pairs across all FIR-related continuous components. Random-pair delta divided by adjacent-pair delta was 1.72 for MAF, 2.17 for |BETA|, 2.29 for |Z|, 2.28 for LP and 1.63 for SE, all with empirical P = 9.99 × 10^-4 (Fig. 7B). Thus, FIR spatial regularity is rooted in local autocorrelation of the underlying statistical variables, not only in discrete profile labels.

F/I/R decomposition showed that each component contributed to same-profile adjacency, but the full FIR8 combination produced the strongest signal. Single-component ratios ranged from 1.33 to 1.44, pairwise combinations increased to approximately 1.92–2.04, and full FIR8 reached 2.98 (Fig. 7C). Distance-stratified analysis further showed that FIR8 enrichment remained above permutation across all adjacent-pair distance bins, decreasing from 2.52 at 0–100 bp to 1.71 beyond 50 kb (Fig. 7D). Physical proximity therefore modulates, but does not fully explain, FIR same-profile adjacency.

Domain-level analyses reinforced this interpretation. Observed FIR domains showed markedly lower within-domain variance in MAF, |BETA| and |Z| than matched random segments, with log2 ratios of -2.58, -1.81 and -2.17, respectively, whereas pairwise F/I/R correlations changed little (Fig. 7E). This indicates that FIR domains are defined mainly by local homogeneity of component values rather than stronger internal correlation among components. Boundary analysis showed modest but significant shifts in local statistical state across FIR-domain boundaries, with observed/permutation transition ratio of 1.048 (empirical P = 0.001996; Fig. 7F). Finally, adjacent-pair modelling showed that a distance-only model explained little of same-FIR8 adjacency, whereas adding frequency, uncertainty, effect, reliability and significance variables increased McFadden R² to 0.5807 (Fig. 7G).

Together, these findings suggest that FIR-domain architecture emerges from coordinated local continuity in allele frequency, effect magnitude and statistical reliability, with domain boundaries marking transitions between local genetic-statistical states.

## 4 Discussion

### 4.1 Principal findings

This study developed FIR-GWAS to test whether GWAS summary statistics contain reproducible local genetic-statistical organization beyond individual association peaks^2,20^. The central finding is that SNPs with similar frequency, impact and reliability profiles are not randomly distributed along genomic coordinates, but show consistent same-profile continuity and form coordinate-contiguous FIR-defined domains. This pattern was first observed in EUR height GWAS, persisted after removing genome-wide significant variants and chromosome-level extreme association signals, and recurred across ancestry-specific height datasets and eight additional human complex traits.

Extending local continuity to domain-level analysis showed that same-profile SNPs form longer and denser blocks than expected under chromosome-stratified permutation. FIR8 domain calling further separated association strength from regional structure: some genome-wide significant regions formed structured multi-SNP domains, whereas others remained isolated, and a subset of non-GWS subthreshold regions showed coordinated domain structure despite not crossing the conventional significance threshold. Biological annotation connected structured domains to regulatory and height-related gene contexts^4,5,11^. Cross-species analyses showed recurrence in Arabidopsis and chicken association datasets, and genetic-statistical decomposition indicated that FIR spatial regularity is rooted in local similarity of allele frequency, effect magnitude and statistical reliability. Together, these findings support FIR-domain architecture as a domain-level layer of GWAS summary-statistic organization.

### 4.2 FIR-domain architecture as a new layer of GWAS summary-statistic organization

Conventional GWAS interpretation is largely organized around single-variant significance, lead SNPs and locus-level association peaks^2,20^. This strategy has identified many trait-associated loci, but it usually treats the surrounding summary-statistic landscape as supporting information rather than as an object of analysis^6^. FIR-GWAS instead treats GWAS summary statistics as a coordinate-ordered genetic-statistical landscape in which each SNP carries a local state defined jointly by allele frequency, effect magnitude and statistical reliability.

The main conceptual advance is the transition from local FIR continuity to FIR-domain architecture. FIR domains are defined as coordinate-contiguous same-profile SNP runs rather than as externally imposed genes, loci, linkage disequilibrium blocks, chromatin domains or fixed-width windows^17,21^. Their enrichment for high-percentile length, SNP count and N50-based metrics under chromosome-stratified permutation indicates that these domains are not expected from random placement of profile labels while preserving chromosome-specific SNP density. In this sense, FIR domains provide a statistical-genomic representation of local GWAS architecture.

FIR domains should not be interpreted as experimentally defined biological domains. They do not directly imply chromatin looping, regulatory activity or causal locus boundaries. Their value is as an intermediate layer between individual SNPs and downstream biological annotation: a reproducible, coordinate-aware and permutation-testable structure derived from GWAS summary statistics. This layer allows GWAS results to be interpreted not only by whether a lead SNP is significant, but also by whether a local genomic segment carries a coordinated frequency–impact–reliability state.

### 4.3 Relationship to conventional GWAS significance and subthreshold signals

A key implication of FIR-domain calling is that genome-wide significance and regional structure are related but separable properties of GWAS signals^20,22^. Conventional GWAS prioritizes variants or loci by whether association statistics exceed a predefined threshold^23^. FIR-domain calling asks a different question: whether neighbouring SNPs share a coordinated frequency–impact–reliability profile and form a domain that is longer or denser than expected under chromosome-stratified permutation.

The distinction is illustrated by GWS structured and GWS non-structured domains. GWS structured domains contain genome-wide significant variants embedded within multi-SNP FIR structure, suggesting regional statistical support. GWS non-structured domains also contain genome-wide significant variants but fail the structured-domain criteria, indicating that a strong individual association peak does not necessarily imply coordinate-contiguous regional structure^22^. FIR-GWAS therefore adds a second axis of interpretation: association strength and regional structure can be evaluated jointly rather than treated as equivalent.

The non-GWS subthreshold structured domains provide the clearest example of the added value of this framework^23^. These regions do not contain variants crossing the conventional genome-wide significance threshold, but they pass the q95 structured-domain definition and show coordinated multi-SNP FIR architecture. They should not be interpreted as confirmed associations or substitutes for genome-wide significant loci^24^. Instead, they represent prioritized subthreshold regions with non-random local statistical structure, suitable for downstream annotation, replication and functional follow-up. Thus, FIR-GWAS complements conventional GWAS by preserving the value of genome-wide significance while exposing domain-level information not visible from lead SNPs alone.

### 4.4 Biological interpretability and candidate-domain prioritization

Although FIR domains are defined from GWAS summary-statistic profiles rather than biological annotations, their interpretive value depends on whether they connect to meaningful genomic context^4,5,11^. In the height analysis, GWS structured domains were enriched for promoter-like, enhancer-like and accessible chromatin annotations and overlapped many curated skeletal-growth and developmental signalling genes^10,11^. In contrast, GWS non-structured domains showed a more isolated statistical profile and were comparatively depleted for several regulatory annotations. This supports the distinction between a significant association peak and a structured regulatory genomic context^20^.

Non-GWS subthreshold structured domains were particularly relevant for candidate prioritization^22,25^. These domains did not cross the conventional genome-wide significance threshold but showed high structure scores, multi-SNP regional organization and regulatory annotation support. They also contained or neighboured subsets of height-related genes from skeletal-growth, developmental signalling, growth-plate, extracellular-matrix and endocrine growth-axis gene sets. Candidate regions near BMPR1B, ADAMTS17, TGFBR2, COMP, WNT10B, LRP5, GLI1 and COL11A1 illustrate how FIR-domain structure can highlight biologically plausible subthreshold regions that would be under-emphasized by a strict single-SNP framework.

These findings support FIR domains as biologically interpretable prioritization units, but not as functional validation^24,26^. Enrichment for cCRE, Roadmap or height-related gene annotations does not establish causal variants, causal genes or regulatory mechanisms^5,26^. Rather, FIR-domain annotation provides a structured statistical framework for ranking regions that combine regional FIR organization, association intensity and biological context. Functional interpretation of prioritized domains will require independent evidence from fine-mapping, molecular QTL colocalization, experimental perturbation or trait-relevant cellular assays^24,26^.

### 4.5 Recurrence across traits, ancestries and species

The recurrence of FIR-domain architecture across analytical contexts argues against a dataset-specific artefact^2,20^. In height GWAS, FIR continuity was first detected in EUR but also appeared across ancestry-specific datasets despite large differences in association-enriched SNP counts and profile composition^27^. EUR-anchored profiling further showed that non-EUR datasets differed in FIR composition, indicating that recurrence across ancestries was not simply caused by identical profile distributions.

The extension to eight human complex traits further supports generality^6,28^. FIR continuity was observed across anthropometric, lipid, metabolic, vascular, pulmonary and behavioural traits, despite substantial variation in the number of LP > 4 SNPs and genome-wide significant variants. At the same time, trait-level FIR signatures were not identical; related traits showed more similar patterns, whereas other traits differed^23^. Thus, FIR-domain architecture appears recurrent across human complex traits, while the detailed organization of frequency, impact and reliability remains phenotype-dependent.

Cross-species analyses extended this pattern beyond human GWAS^12^. FIR-domain architecture recurred in Arabidopsis flowering-time traits and chicken growth or production traits, using trait-specific top-fraction association sets and chromosome-stratified permutation backgrounds^12,29^. This recurrence should be interpreted as statistical rather than biological: it does not imply that human height, Arabidopsis flowering time and chicken growth traits share conserved mechanisms. Instead, it suggests that association-summary landscapes from diverse systems can exhibit local frequency–impact–reliability organization when analysed through a coordinate-based framework.

### 4.6 Genetic-statistical basis of FIR spatial regularity

The genetic-statistical decomposition indicates that FIR spatial regularity is rooted in local continuity of the continuous variables underlying the FIR profiles^4,6^. Adjacent SNPs were more similar than chromosome-matched random SNP pairs across MAF, effect magnitude, statistical reliability, association strength and standard error^7,16^. This shows that FIR-domain architecture is not merely an artefact of discretizing SNPs into profile labels, but reflects local autocorrelation in GWAS-derived statistical variables^6^.

Frequency, impact and reliability each contributed to same-profile adjacency, but no single component fully explained the observed pattern. Single-component profiles showed enrichment, pairwise combinations strengthened the signal, and the full FIR8 profile produced the strongest same-profile enrichment. Distance also contributed, with stronger enrichment among closer adjacent SNPs, but distance-stratified analyses and adjacent-pair modelling showed that physical proximity alone was insufficient^7,21^. The architecture therefore depends on the joint spatial behaviour of allele frequency, effect magnitude and reliability rather than on any single component or distance alone.

At the domain level, observed FIR domains were characterized mainly by within-domain homogeneity of MAF, |BETA| and |Z| rather than by substantially stronger internal correlations among FIR components^4^. Boundary analyses further showed transitions in local genetic-statistical state at FIR-domain limits. Together, these findings support a model in which FIR domains represent locally homogeneous statistical segments separated by shifts in frequency–impact–reliability state. This provides a genetic-statistical rationale for why FIR-GWAS can detect reproducible same-profile domains while remaining a summary-statistic framework rather than a direct model of molecular function^6^.

### 4.7 Limitations and conclusions

Several limitations should be considered. FIR-GWAS is based on GWAS summary statistics rather than individual-level genotype and phenotype data, so it cannot directly model haplotypes, individual-level prediction or causal genotype–phenotype relationships^6,20^. Although the permutation procedures preserve genomic coordinates, chromosome structure, SNP density or local window structure depending on the analysis, they do not provide a full causal decomposition of linkage disequilibrium, imputation structure, marker density or SNP ascertainment^7^. FIR-domain architecture should therefore be interpreted as summary-statistic organization, not as a direct molecular mechanism.

FIR profiles and structured-domain calls also depend on analytical choices, including LP threshold, FIR split strategy, percentile cutoff and minimum-SNP requirement. Robustness analyses showed that the main signal persisted across alternative thresholds, null models, structured-domain definitions, regional exclusions and FIR splitting strategies, but the exact number and boundaries of called domains remain framework-dependent. Biological annotation was used for interpretation and prioritization, not validation, and cross-species recurrence should be interpreted as recurrence of statistical architecture rather than conserved biology across species^26^.

In conclusion, FIR-GWAS identifies a domain-level layer of local genetic-statistical organization in GWAS summary statistics. By integrating allele frequency, effect magnitude and statistical reliability, the framework detects same-profile continuity, converts it into scoreable genomic domains and separates regional structure from single-variant significance. FIR-GWAS complements conventional lead-SNP analysis by distinguishing GWS structured regions, isolated GWS peaks and non-GWS subthreshold structured domains, providing a summary-statistic framework for detecting, scoring and interpreting local genomic architecture beyond conventional GWAS significance.

## 5 Conclusion

We show that GWAS summary statistics contain a previously unrecognized layer of local genetic–statistical organization structured along genomic coordinates. Using FIR-GWAS, we demonstrate that allele frequency, effect magnitude and statistical reliability jointly define coordinate-contiguous domains that are enriched beyond null expectations across multiple traits and species.

These FIR-domain structures persist beyond genome-wide significant loci and reveal a domain-level organization within GWAS summary statistics. This framework suggests that GWAS signals are better interpreted as structured genetic–statistical landscapes rather than independent variant-level associations, providing a complementary layer to conventional locus-based analysis.

## Author Contributions

Hongdong Hao, Dian Chen, and Xinyu Zhang performed data processing and statistical analyses. Hui Xue and Tianyu Meng conceived and supervised the study. Hongdong Hao and Dian Chen contributed to FIR-GWAS implementation and figure generation. Xinyu Zhang contributed to GWAS data curation and methodological support. Hui Xue and Tianyu Meng provided clinical interpretation and critical revision of the manuscript. All authors contributed to manuscript writing and approved the final version.

## Generative AI Statement

During the preparation of this manuscript, the authors used OpenAI’s ChatGPT solely as a language editing and proofreading aid. All scientific content, analyses, interpretations, and conclusions were developed and verified by the authors. The authors take full responsibility for the integrity and accuracy of the manuscript.

## References

1. Manolio, T.A. et al. Finding the missing heritability of complex diseases. Nature 461, 747–53 (2009).

2. Visscher, P.M. et al. 10 Years of GWAS Discovery: Biology, Function, and Translation. Am J Hum Genet 101, 5–22 (2017).

3. Purcell, S. et al. PLINK: a tool set for whole-genome association and population-based linkage analyses. Am J Hum Genet 81, 559–75 (2007).

4. Finucane, H.K. et al. Partitioning heritability by functional annotation using genome-wide association summary statistics. Nat Genet 47, 1228–35 (2015).

5. Maurano, M.T. et al. Systematic localization of common disease-associated variation in regulatory DNA. Science 337, 1190–5 (2012).

6. Bulik-Sullivan, B.K. et al. LD Score regression distinguishes confounding from polygenicity in genome-wide association studies. Nat Genet 47, 291–5 (2015).

7. Gabriel, S.B. et al. The structure of haplotype blocks in the human genome. Science 296, 2225–9 (2002).

8. Yengo, L. et al. A saturated map of common genetic variants associated with human height. Nature 610, 704–712 (2022).

9. MacArthur, J. et al. The new NHGRI-EBI Catalog of published genome-wide association studies (GWAS Catalog). Nucleic Acids Res 45, D896–d901 (2017).

10. An integrated encyclopedia of DNA elements in the human genome. Nature 489, 57–74 (2012).

11. Kundaje, A. et al. Integrative analysis of 111 reference human epigenomes. Nature 518, 317–30 (2015).

12. Atwell, S. et al. Genome-wide association study of 107 phenotypes in Arabidopsis thaliana inbred lines. Nature 465, 627–31 (2010).

13. Aranzana, M.J. et al. Genome-wide association mapping in Arabidopsis identifies previously known flowering time and pathogen resistance genes. PLoS Genet 1, e60 (2005).

14. Pickrell, J.K. Joint analysis of functional genomic data and genome-wide association studies of 18 human traits. Am J Hum Genet 94, 559–73 (2014).

15. Chang, C.C. et al. Second-generation PLINK: rising to the challenge of larger and richer datasets. Gigascience 4, 7 (2015).

16. Winkler, A.M., Ridgway, G.R., Webster, M.A., Smith, S.M. & Nichols, T.E. Permutation inference for the general linear model. Neuroimage 92, 381–97 (2014).

17. Ernst, J. & Kellis, M. ChromHMM: automating chromatin-state discovery and characterization. Nat Methods 9, 215–6 (2012).

18. Harrow, J. et al. GENCODE: the reference human genome annotation for The ENCODE Project. Genome Res 22, 1760–74 (2012).

19. Hughes, G., Choudhury, R.A. & McRoberts, N. Summary Measures of Predictive Power Associated with Logistic Regression Models of Disease Risk. Phytopathology 109, 712–715 (2019).

20. Dehghan, A. Genome-Wide Association Studies. Methods Mol Biol 1793, 37–49 (2018).

21. Berisa, T. & Pickrell, J.K. Approximately independent linkage disequilibrium blocks in human populations. Bioinformatics 32, 283–5 (2016).

22. Eichler, E.E. et al. Missing heritability and strategies for finding the underlying causes of complex disease. Nat Rev Genet 11, 446–50 (2010).

23. Biological insights from 108 schizophrenia-associated genetic loci. Nature 511, 421–7 (2014).

24. Reed, E. et al. A guide to genome-wide association analysis and post-analytic interrogation. Stat Med 34, 3769–92 (2015).

25. Yang, J. et al. Common SNPs explain a large proportion of the heritability for human height. Nat Genet 42, 565–9 (2010).

26. Wen, X., Pique-Regi, R. & Luca, F. Integrating molecular QTL data into genome-wide genetic association analysis: Probabilistic assessment of enrichment and colocalization. PLoS Genet 13, e1006646 (2017).

27. Auton, A. et al. A global reference for human genetic variation. Nature 526, 68–74 (2015).

28. Bulik-Sullivan, B. et al. An atlas of genetic correlations across human diseases and traits. Nat Genet 47, 1236–41 (2015).

29. Rubin, C.J. et al. Whole-genome resequencing reveals loci under selection during chicken domestication. Nature 464, 587–91 (2010).

